# Evaluating research-grade and commercial SFDI platforms for burn severity assessment and feature reduction

**DOI:** 10.64898/2026.07.27.741081

**Authors:** Christopher A. Campbell, Gordon T. Kennedy, Alberto Martín-Pérez, Theresa L. Chin, Victor Joe, Robert J. Christy, Anthony J. Durkin

## Abstract

Spatial frequency domain imaging (SFDI) has demonstrated the ability to provide early, quantitative assessment of burn wound severity. Previous studies using a research-grade SFDI platform showed that high-dimensional datasets incorporating multiple spatial frequencies and wavelengths can predict healing outcomes in controlled porcine models of graded burns. To assess the impact of reduced measurement dimensionality on diagnostic performance, we compared classifications derived from a research-grade SFDI system (Reflect RS) with those obtained using a simplified commercial SFDI platform (Clarifi RS). Both systems were used to image graded burns 24 hours after injury. Pixel-level classifiers were trained using regions defined by 28-day healing outcomes, and models based on the full Reflect dataset were compared with classifiers generated from reduced Reflect feature sets and datasets designed to mimic Clarifi acquisition features. Performance was evaluated using leave-one-subject-out validation. The full Reflect dataset achieved the highest classification performance, with mean F1 scores approaching 0.88. Although reducing the number of measured features decreased classification accuracy, simplified models and Clarifi-based datasets maintained F1 scores greater than 0.8 for binary classification. These findings indicate that dimensionality reduction produces a measurable but manageable loss in performance and support the development of clinically practical, application-specific SFDI systems for burn assessment.

## 1 Introduction

### 1.1 Burn Severity Classification

Approximately 400,000 people are treated for burn wounds in the US annually^1^. Management in the first 72 hours following a burn is critical, as early intervention is linked to improved outcomes^2,3^. However, research has shown that physicians are somewhere between 60% and 75% accurate in assessing burn severity, particularly in differentiating superficial partial thickness burns (which typically can heal with appropriate wound care) from deep partial thickness burns (which are often managed with surgical intervention) in the first 72 hours^4,5^. An accurate and quantitative measure of early burn severity would enable physicians to provide earlier management of wound healing, leading to better functional and cosmetic outcomes for patients and potentially allowing for reduced inpatient duration. We have previously reported on the recent development of objective technologies for non-invasive burn severity assessment^6,7^. While technologies like hyperspectral, laser speckle, and laser Doppler imaging are notable in this space, our group specializes in spatial frequency domain imaging (SFDI). We have actively developed and characterized this quantitative, noninvasive, wide-field modality specifically with the goal to improve burn severity assessment.

### 1.2 Spatial Frequency Domain Imaging (SFDI)

SFDI is a depth-sensitive, quantitative widefield imaging technology useful for investigating *in vivo* tissues, including skin. The details of SFDI as practiced by our group have been described in the literature^8,9^. Briefly, in a common configuration, structured illumination in the form of sinusoidal irradiance patterns is projected onto a target tissue of interest at two or more spatial frequencies and three phase offsets per frequency, at multiple wavelengths in the visible to near infrared region of the electromagnetic spectrum (e.g., 470-850 nm). The three images for a given spatial frequency and optical wavelength are demodulated and calibrated against images of a tissue-mimicking phantom with known optical properties to obtain the AC amplitudes of the modulation envelopes at every pixel.

These “diffuse reflectance” maps, representing the modulation transfer function of the target tissue, are either fitted by regression to a forward model of light transport or compared to a lookup table to find the optical properties (namely, wavelength dependent absorption and reduced scattering coefficients) of the interrogated tissue. The absorption spectrum can be further decomposed into chromophore concentration maps of such species as oxy- and deoxyhemoglobin, water, lipids, and melanin^10^. Alternatively, the calibrated reflectance can be used, without invoking a model of light transport, within the context of a machine learning framework^11^. It is this scenario that we have previously found promising within the context of burn severity assessment. Here we will apply a similar approach to investigate and compare the performance of a simplified SFDI device (Clarifi RS) to the fully featured research grade SFDI device (Reflect RS).

A major advantage of SFDI over conventional widefield imaging is that its detection depth varies based on the projected spatial frequency and wavelength used used.^12^ While methods to analyze this interplay are still evolving, we have found that this depth dependence offers greater predictive accuracy than standard multispectral imaging without patterned illumination, as demonstrated by Rowland et al^9^.

### 1.3 Objective: SFDI for Burn Assessment

Our group has previously demonstrated that SFDI-derived features correlate with burn severity in porcine models of graded burn severity and can classify healing outcomes within 24 hours of burn injury^9,13,14^. The study described in Rowland *et al* relied on a research-grade platform SFDI device (Reflect RS; Modulim, Irvine CA) with dense sampling across wavelength (eight LEDs spanning 471 nm to 851 nm) and spatial frequency (0/mm to 0.2/mm in 5 equal steps) domains^9^. While effective, this configuration is not optimized for efficient, clinically relevant burn related workflow, particularly for challenging environments relevant to triage of military trauma or for mass casualty scenarios. More recently, regression-based analysis demonstrated that calibrated SFDI reflectance can predict collagen denaturation depth and healing outcome while also identifying spectral and spatial-frequency combinations that contribute strongly to burn severity assessment^15^. The analyses presented in Phan et. al. were performed entirely within a research-grade acquisition framework.

The Clarifi (Modulim, Irvine CA) system represents a clinically deployable alternative (Fig. 1) , with a smaller footprint, enhanced LED brightness (reducing motion artifact related to respiration and ambient light) reduced spectral and spatial sampling, rapid acquisition, and integrated processing^16^. This simplification reduces system complexity but also reduces available measurement information.

**Figure 1.**
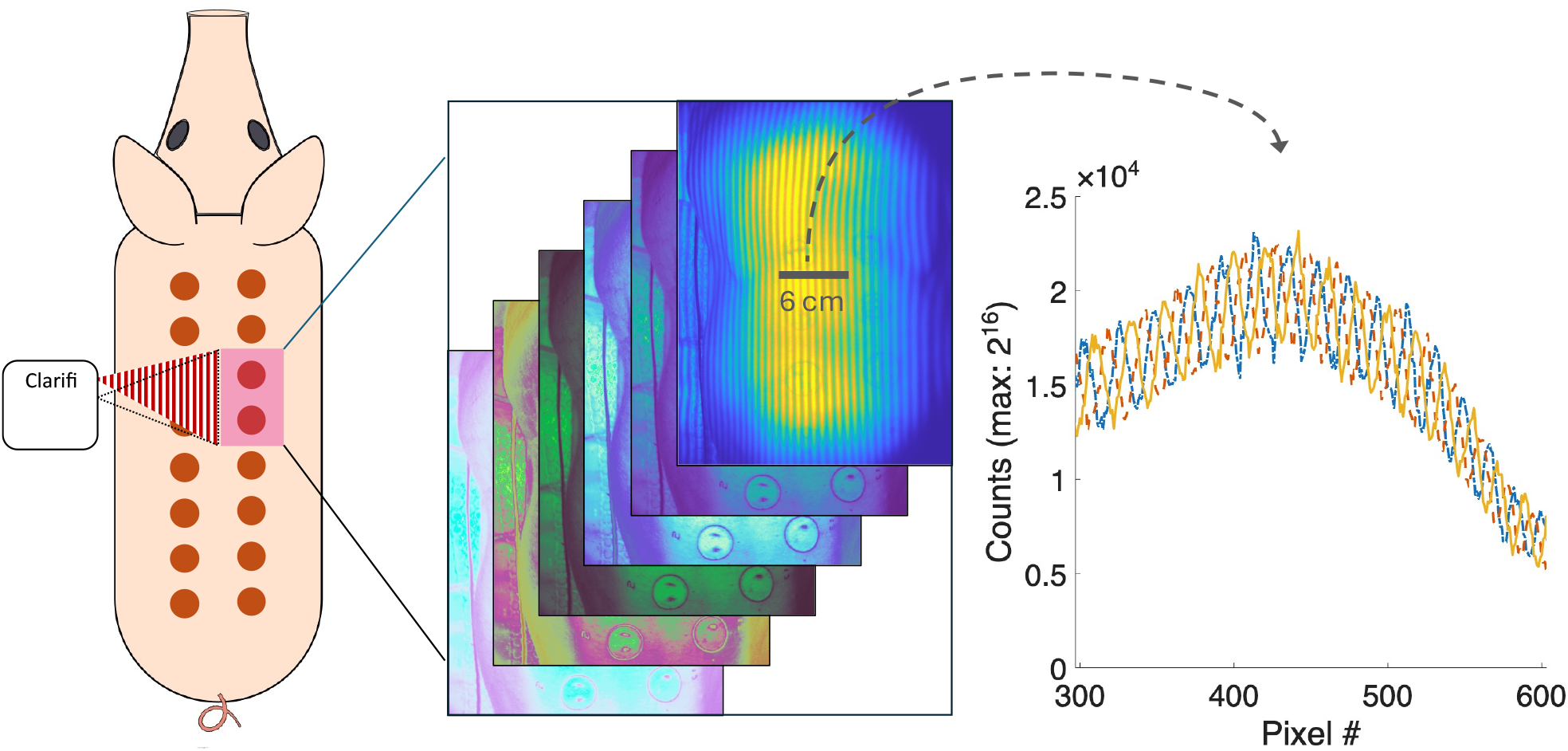
The Clarifi imager employs a series of flood-illuminating LEDs for unmodulated, or planar, images (center wavelengths ∼ 470, 525, 620, 730, and 850 nm), alongside a single projected pattern at 850 nm (spatial frequency 0.12 mm^-1^), generated with a sinusoidal mask and physically shifted to generate three phase offsets (0, 2π/3 and 4π/3). The system is positioned at its specified working distance from the animal, with two burns centered in the field of view. The inset on the right shows a small cross section of the three modulated images, which are used to find the modulation amplitude.

The present work evaluates the impact of this reduction on classification performance using a porcine model of graded burns similar to that previously presented^9^. We compare burn severity prediction using models trained on the Reflect RS datasets to Clarifi-trained models to isolate the effect of feature dimensionality on burn classification. Specifically, we train models on an expanded set of subjects and apply modern analysis methods to obtain a more robust predictor of model performance. We then perform the same analysis using data from an “off the shelf” version of Clarifi. The performance of models trained for use with the new instrument, using six features, is compared to the performance of the original, 40-parameter instrument. For a robust analysis, we use Leave-one-subject-out (LOSO) evaluation when assessing our models^17^.

## 2 Materials and Methods

### 2.1 Instrumentation

The Reflect RS (known as “OxImager RS” prior to a name change by the manufacturer), is a research-grade device (circa 2014, Modulim Inc., Irvine), with up to ten modulated wavelengths configurable for a wide range of spatial modulation frequencies. Under the configuration used for this work, eight wavelengths were collected at a minimum of five spatial frequencies (0/mm to 0.2/mm in 0.05/mm increments), for a total of 40 available spatial frequency/optical wavelength combinations, or “features,” at every pixel. Because the Reflect RS (distinct from an updated Modulim offering, the Reflect 2) has an intermittent timing issue that results in occasional demodulation artifacts, we collect three repetitions of each measurement sequence, resulting in an acquisition time of roughly ninety seconds (this also enables us to disregard acquisitions that have gross movement artifacts such as those resulting from respiration). Processing is performed on a desktop computer using software provided by Modulim. A typical data cube (after processing, but before cropping, labeling, and training) example is shown in Figure 2(a), with a Clarifi data cube shown in 2(b).

**Figure 2.**
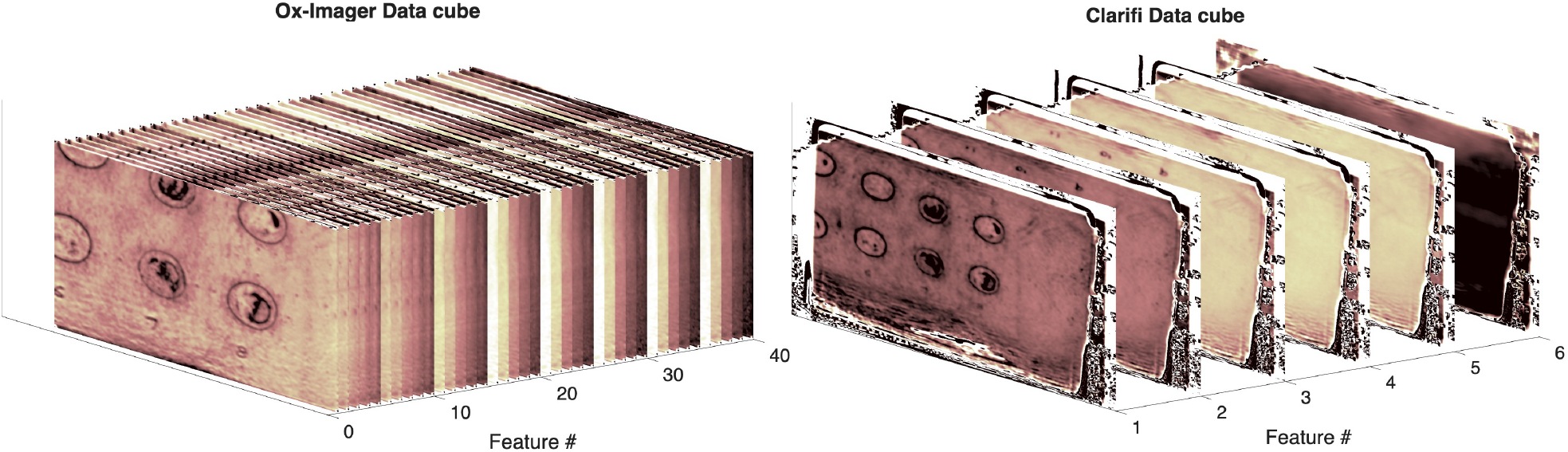
Calibrated reflectance datacubes: Left: Reflect RS data, comprising 8 wavelengths, each with 5 different spatial frequencies. Right: Clarifi datacube. Five planar images (0 mm^-1^), plus the modulated image at 850 nm.

Like the clinical version of the Reflect RS (known as the OxImager CS), the Clarifi imager is FDA-cleared for tissue oximetry, with features optimized for assessing perfusion in the legs and feet with a goal of identification of skin at risk for formation of ulcers in diabetic patients having compromised peripheral vasculature. Specific optimizations of the Clarifi over the Reflect RS include improved source brightness, fewer wavelengths, fewer spatial frequencies, achieving an acquisition time of less than ten seconds and a signal-to-noise ratio improvement of Clarifi under ambient lights. The system, in its off-the-shelf clinical usage configuration for wide-field oximetry, acquires image data and processes it on the attached tablet for an additional 7-8 seconds, after which interactive maps of oxygen saturation and other metrics are displayed. Modulim has slightly modified the configuration used in our investigation in terms of control software (allowing a “research mode”). This modified system is labeled “Clarifi RS”. While the clinical Clarifi system is set up for imaging the sole of a foot held out in front of a patient, the cart used for the “RS” systems affords more degrees of freedom, enabling ease of use both for porcine experiments and in the burn unit with clinical burn data collection. Where the images are typically processed on device with the clinical configuration, the images collected within the context of our burn studies were processed on a separate desktop, as with the Reflect RS.

A schematic of the Clarifi imaging concept appears in Fig 1. Not shown is the calibration step common to SFDI systems^8^, which must be performed at least once per day for the SFDI devices used here. The data acquired in this step consists of a complete image stack of a calibration standard (tissue simulating phantom having known optical properties, and similar to the target tissue of interest), to be used for determining the system response when processing data. Representative images collected by the Clarifi system, which are subsequently cropped to evaluate each of two burns in the center of the image, are compared with images from the Reflect RS system in Fig 3.

**Figure 3.**
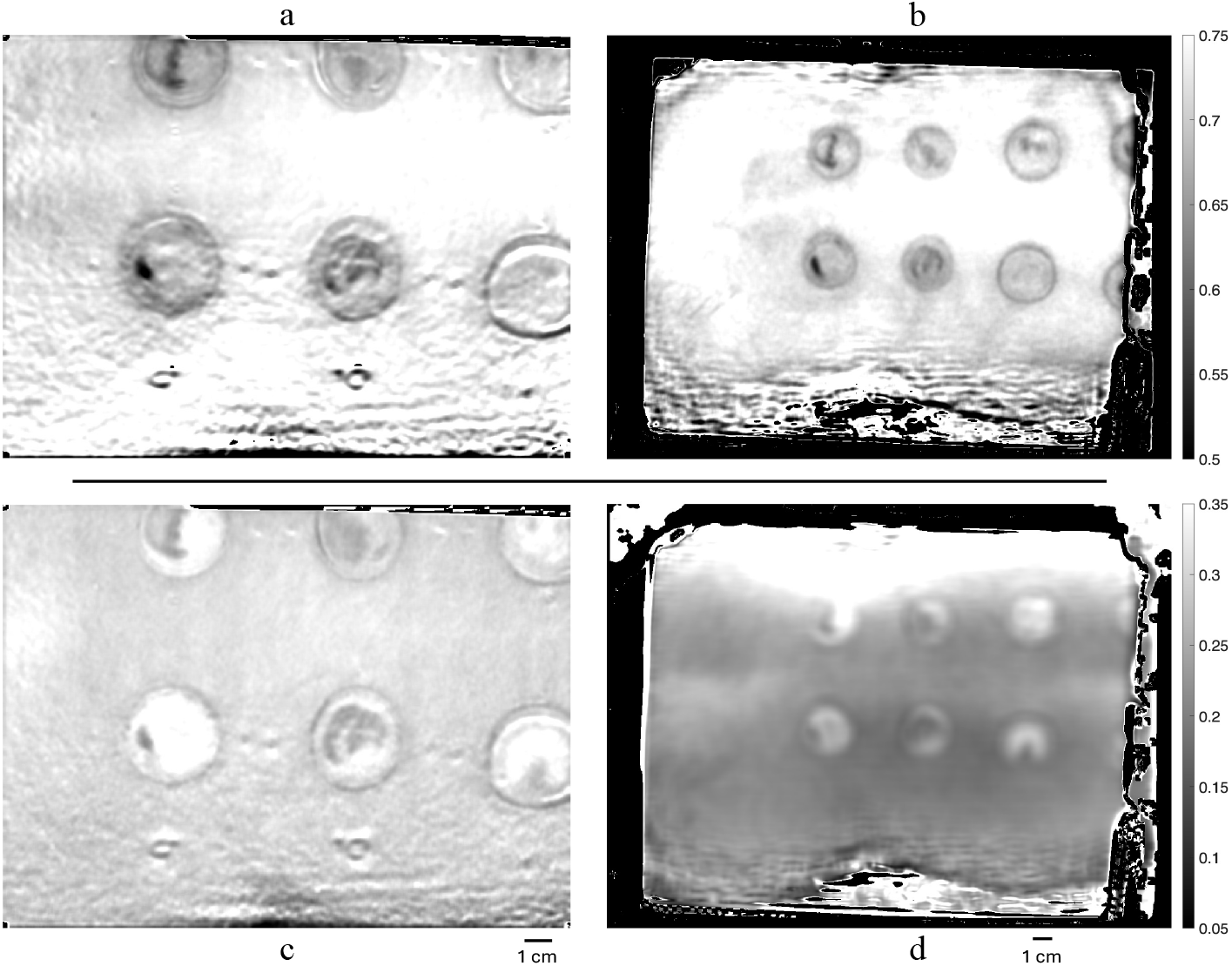
Top row: Calibrated planar reflectance for the Reflect RS (left) and Clarifi RS (right). Some modulation artifacts are visible in both systems, which may be due to the profilometry algorithm struggling to account for steep angles, and/or breathing motion. Bottom row: Calibrated AC reflectance at 850 nm. 0 for the Reflect RS (left) and Clarifi RS (right). The range of values displayed in the calibrated reflectance of the modulated image is substantially smaller than the others, as it represents the modulation amplitude of the projected patterns.

### 2.2 Porcine Model

The porcine models used for these investigations of burn wound imaging provide a controlled environment in which to develop our methods. The specifics have been detailed before^13^. Briefly, sixteen 3 cm burn wounds, were created, using a custom <u>custom-</u>made burn tool, with a range of severities on each pig (N=4, Yorkshire/Landrace hybrid, 45-65 kg) under an IACUC-approved protocol (UCI # AUP-21-014 and AUP-24-056). All animal research was conducted in compliance with the Animal Welfare Act, the implementing Animal Welfare regulations, and the principles of the Guide for the Care and Use of Laboratory Animals. The facility where the research was conducted is fully accredited by the AAALAC International.

Images were framed to contain two burns each and were acquired sequentially using the SFDI devices approximately 1 hour after administration of the burns, as well as on days 1, 3, 7, 14, 21, and 28 post-burn. Color photographs, providing a record of clinical appearance, were also acquired at each time point with a 14-megapixel digital handheld camera (Sony Alpha NEX-3). Photo documentation of clinical appearance using a high quality digital camera with consistent illumination has been a standard practice in the group for these investigations^9^ because the quality of the RGB camera included in each SFDI device varies and those images typically have poor spatial resolution. In this work, photographs were used as a visual record of final clinical healing outcomes as has been done previously^9^. These are used to label SFDI data for model training, as well as for qualitative evaluation of how the classification models performed. Only the SFDI data from day 1 and color photos from day 28 are used for the analysis presented here.

### 2.3 Data Processing and Labeling

Raw SFDI images were processed on an external PC using proprietary software provided for each system by Modulim. There is a box filter applied to Clarifi data to remove demodulation artifacts resulting from the system hardware design. To balance image distortion and effective resolution, enhancing small features necessary for analyzing heterogeneous 3 cm burns, we reduced the box filter window size from 25 pixels (8 mm) to 15 pixels (5 mm). The resulting resolution enhancement is shown in Fig. 4. Reflect RS data were processed using equivalent demodulation parameters without box filtering.

**Figure 4.**
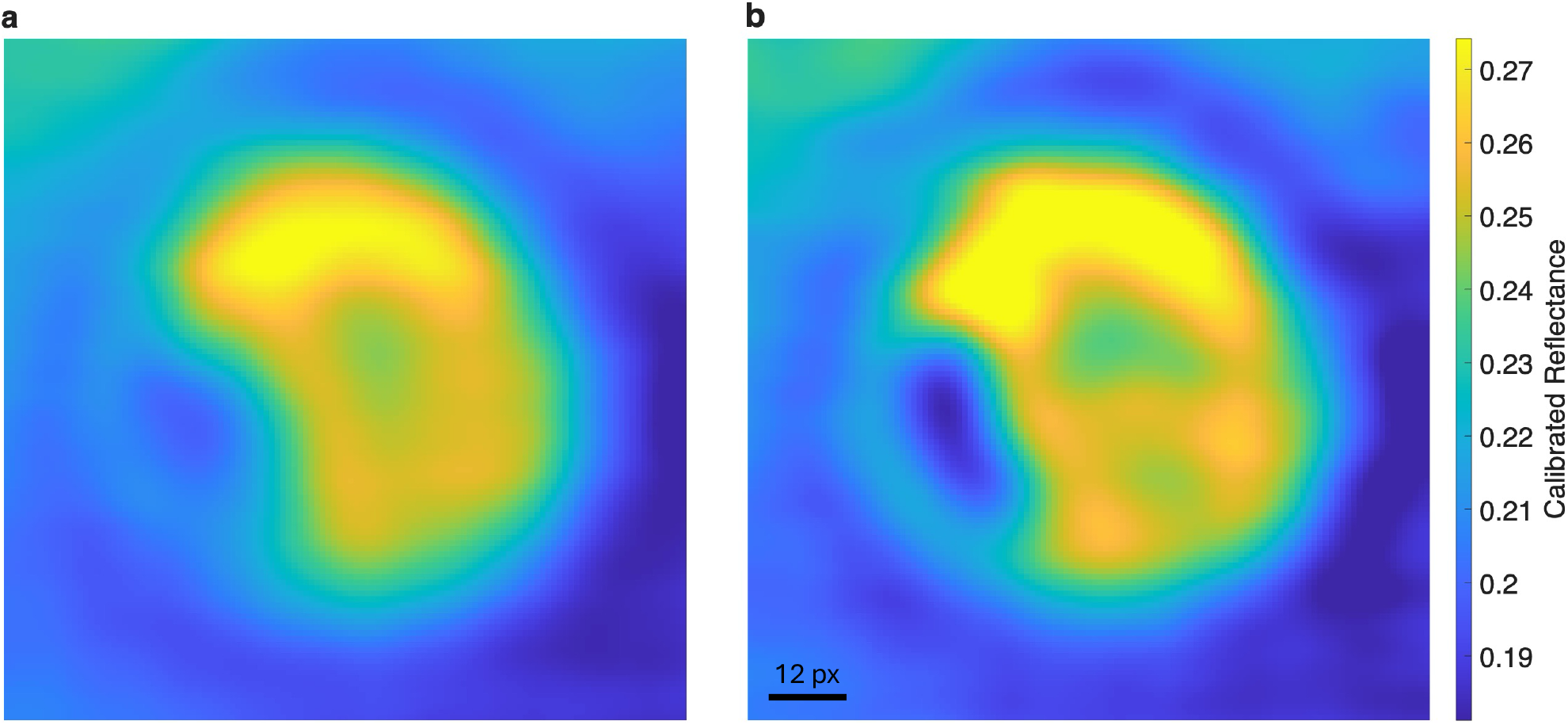
Demodulated images cropped to a burn, with original (**a**) and reduced (**b**) smoothing, showing a resolution advantage at the expense of increased modulation artifacts.

The present labeling procedure is an adaptation of a process set forth in our previous work training a machine learning based classifier^9^. In that work, authors selected five-by-five pixel squares representing four categories from regions based on their appearance in day 28 photographs, calculating mean reflectance values for each square, at each wavelength and spatial frequency. In the present work, two burn researchers separately examined the photographs of each burn from day 28, labeling regions that had re-epithelialized, resulting in fresh healthy skin, and regions that had not healed. Day 28 was chosen as a reference point in time wherein the state of the tissue has revealed whether it should have been grafted (scarred, suggesting a DPT or FT burn severity) or healed with conventional wound care (suggesting SPT burn severity). While not ideal for most studies, this approach to choice of “ground truth” was demonstrated to be relatively robust in our original machine learning feasibility study of the Reflect RS^11^. Where there was consensus, the drawn regions of interest (ROIs) were reproduced on the day 1 Reflect RS and Clarifi RS images in MATLAB. Skin pixels surrounding the burn sites were also conservatively labeled, requiring no specialized knowledge to define. These skin pixels were included in training and evaluating a “trinary” model (skin/viable/nonviable) for each system and excluded for a “binary” model differentiating only between viable and nonviable burn tissue. Landmarks in the SFDI data were identified in the color photographs to guide the ROI selection, based on the first author’s best judgment. Where burns had regions of mixed severity or no consensus, ROIs were drawn conservatively to avoid cross contaminating the training data.

Four pigs, with a total of 64 burns, are used for this analysis, resulting in at least 9,500 individual data points (pixels) from each of the three categories. Where previously a “hyperperfused periphery” classification was included, the authors opted not to do that here. The periphery is only 5-15 pixels across, depending on how its borders are determined. If the same conservative ROI selection method were accurately performed on the periphery, the resulting set of pixels available for training and testing would be quite small. Fig 5 shows a sample of the resulting ROIs.

**Figure 5.**
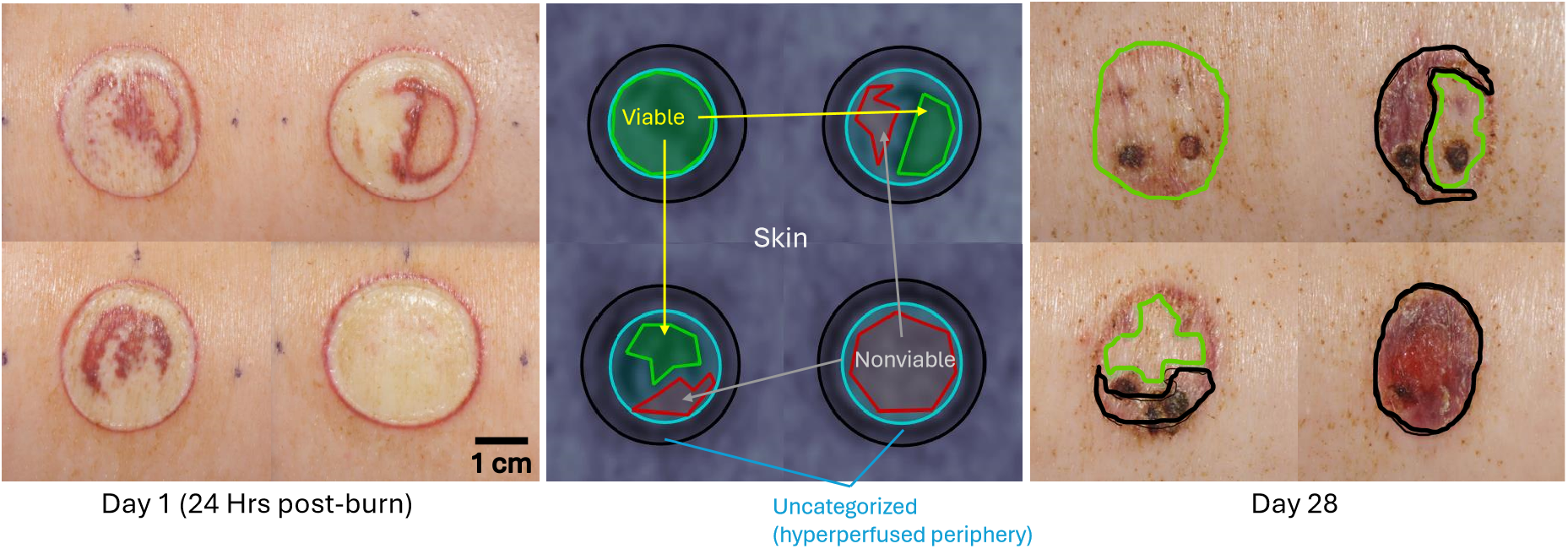
ROI selection. Digital RGB Photographs (no polarizers employed) of representative burns are shown, with Day 1 post-burn on the left, and Day 28 on the right. Burn images from day 28 were classified by researchers into two categories: “Viable” and “Nonviable.” Where the researchers’ assessments agreed, ROIs were selected for model training on Day 1 data (center). For all burns, two circles were drawn (cyan and black), forming an annulus covering the hyperperfused periphery on each burn. These pixels were not labeled, due to uncertainty about which label should be applied. Green polygons were then drawn to outline “viable” pixels, red polygons for “nonviable”, and pixels outside the outer circle were labeled “Skin”. Four biopsy wounds are evident in each burn on Day 28, but were not present in the Day 1 training data.

Every pixel in the Clarifi RS image data was formatted as a vector containing six features: a calibrated reflectance value from each of the five planar images, and the calibrated modulation amplitude (demodulated image) from the 850 nm SFDI images. The Reflect RS values used were all demodulated images, which in the case of the zero-frequency (0/mm) image is equivalent to a planar image. From the labeled pixels, an equal number of random pixels from each category were selected to train a cubic support vector machine (SVM) model.

### 2.4 Evaluating the Classifiers

The trained classifiers were evaluated for accuracy on three bases: The 10-fold cross-validation accuracy reported when the model was trained (included as supplementary material); LOSO performance in the form of an F1 Score for all labeled pixels on each subject; and a qualitative visual assessment of fully classified maps.

K-fold cross-validation is a method of assessing a model’s robustness by systematically training the model on 1−(1*/K*) of the labeled training data and testing the trained model on the remainder. The results reported in the included confusion charts are averaged across the ten folds. This was the method used in our previous work^11^, and we include the results to facilitate comparison with that work.

LOSO evaluation separates the data into training sets consisting of images from three pigs, and testing sets containing all burns from the remaining pig, generating statistics for each of the four testing sets. While K-fold cross-validation is useful for estimating the effectiveness of a model, LOSO evaluation provides a stricter prediction of how the model would perform on a completely new animal subject of similar breed and size—useful for this case where we have a large number of burns but a limited number of subjects. With “Nonviable” pixels deemed “positive” for this work, and “viable” and “skin” forming the set of “negative” pixels, the F1 score evaluates the model’s ability to correctly identify pixels that need intervention.

Finally, a visual assessment of classified burn maps, including regions that were not previously categorized, may provide additional insight into the model’s behavior. These maps will include some unburned skin for the trinary models. In the case of the binary models, the skin is masked out for ease of understanding.

### 2.5 Code Availability

All code (other than Modulim acquisition and processing software) used for this manuscript was written in MATLAB (MathWorks, Natick, MA). Spider plots were constructed using a function from the MathWorks file exchange (https://github.com/NewGuy012/spider_plot/releases/tag/21.0). The finalized code is collocated with the data set at the Dryad link posted in the “Data Availability” section.

## 3 Results

Tables 1 and 2 show the number of pixels available per category for each pig. All categories exceed the 7,500 points used per pig per category used to train the models, but “viable” regions had the fewest pixels available to select from.

**Table 1.**
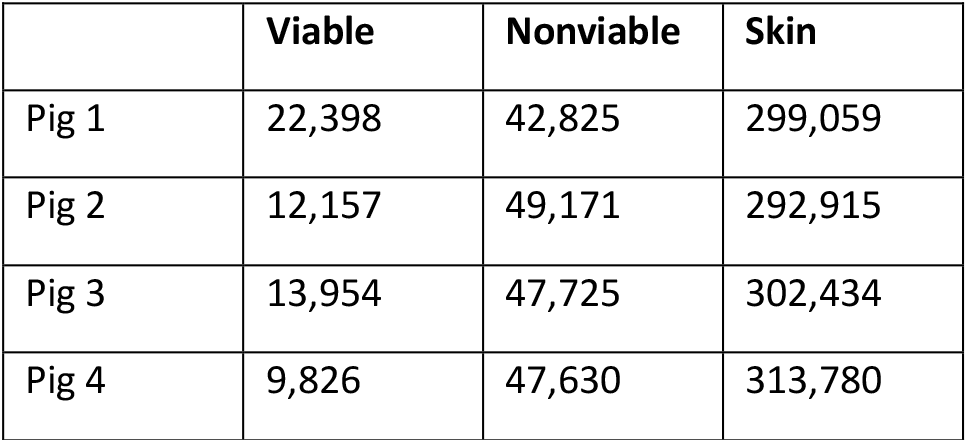
Pixels available in Clarifi data set.

**Table 2.**
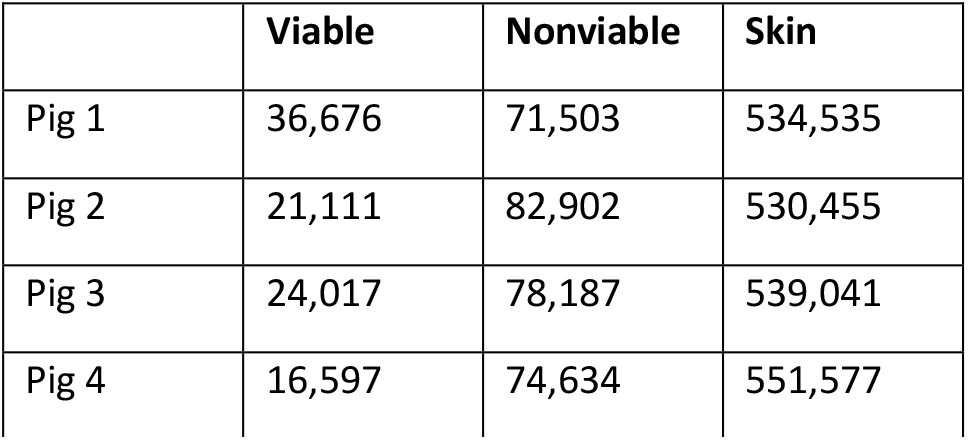
Pixels available in Reflect RS data set.

|  | Viable | Nonviable | Skin |
| --- | --- | --- | --- |
| Pig 1 | 36,676 | 71,503 | 534,535 |
| Pig 2 | 21,111 | 82,902 | 530,455 |
| Pig 3 | 24,017 | 78,187 | 539,041 |
| Pig 4 | 16,597 | 74,634 | 551,577 |

F1 scores and accuracy values for each LOSO round with the trinary model are shown in Figure 6. The Reflect RS trinary models perform according to the expectation set by Rowland et al^11^, with an F1 score and accuracy exceeding 0.8 for every subject (mean: 0.89 for each). Clarifi RS scores are less consistent, with particularly low scores in the case of Pig 4, giving a mean of 0.75. In Figure 7, the classification has been reduced to two categories, “viable” and “nonviable.” This improves the scores of the Clarifi RS classifier to perform similarly well to the full Reflect RS data set, with a mean F1 score of 85 and accuracy of 0.88.

**Figure 6.**
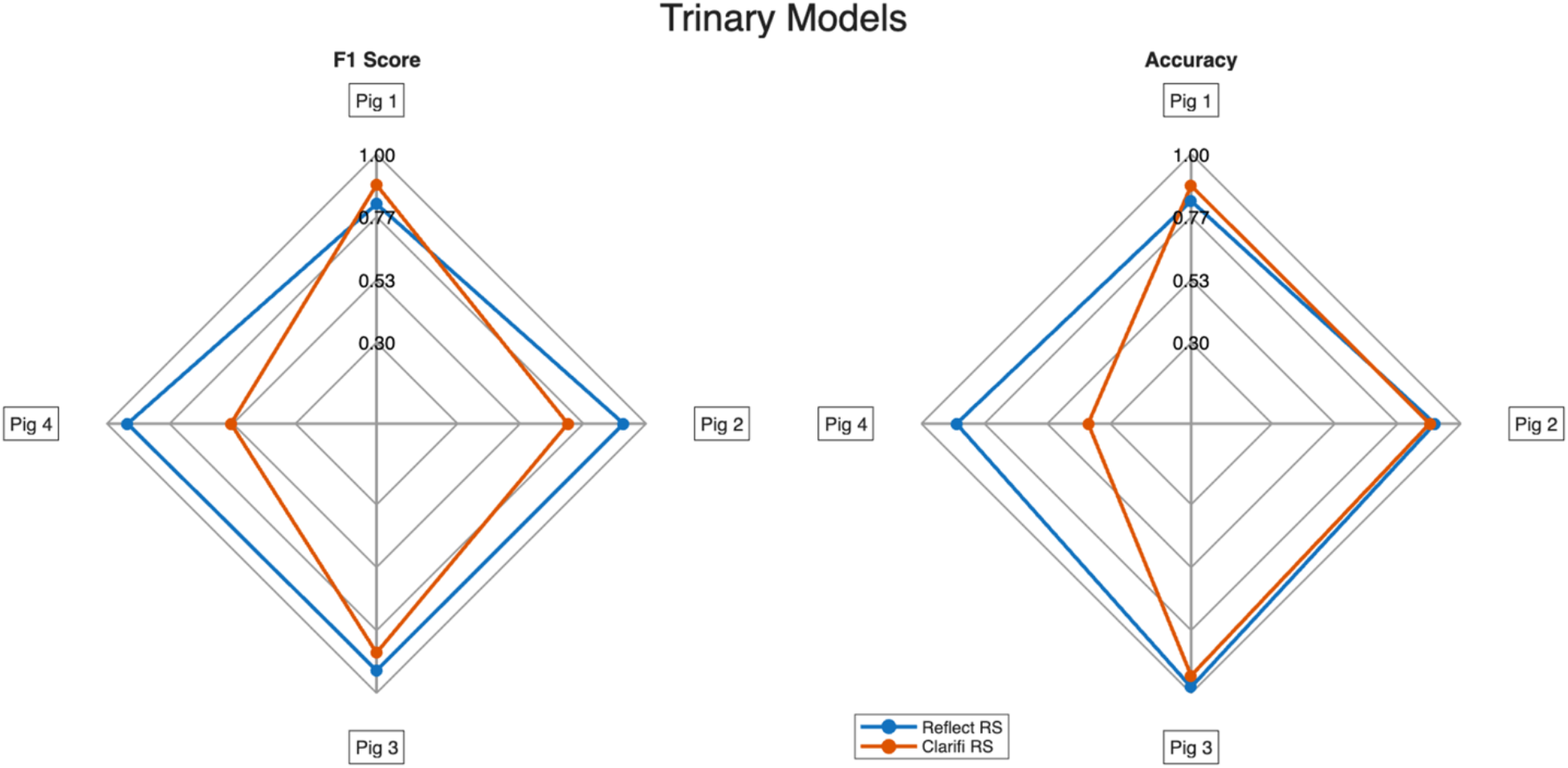
Performance assessment of trinary classifiers. Left: LOSO F1 scores for each pig. Right: LOSO Accuracy in detecting nonviable pixels for each pig. All spider plots were generated using spider_plot() (GitHub: https://github.com/NewGuy012/spider_plot/releases/tag/21.0, accessed July 8, 2026).

**Figure 7.**
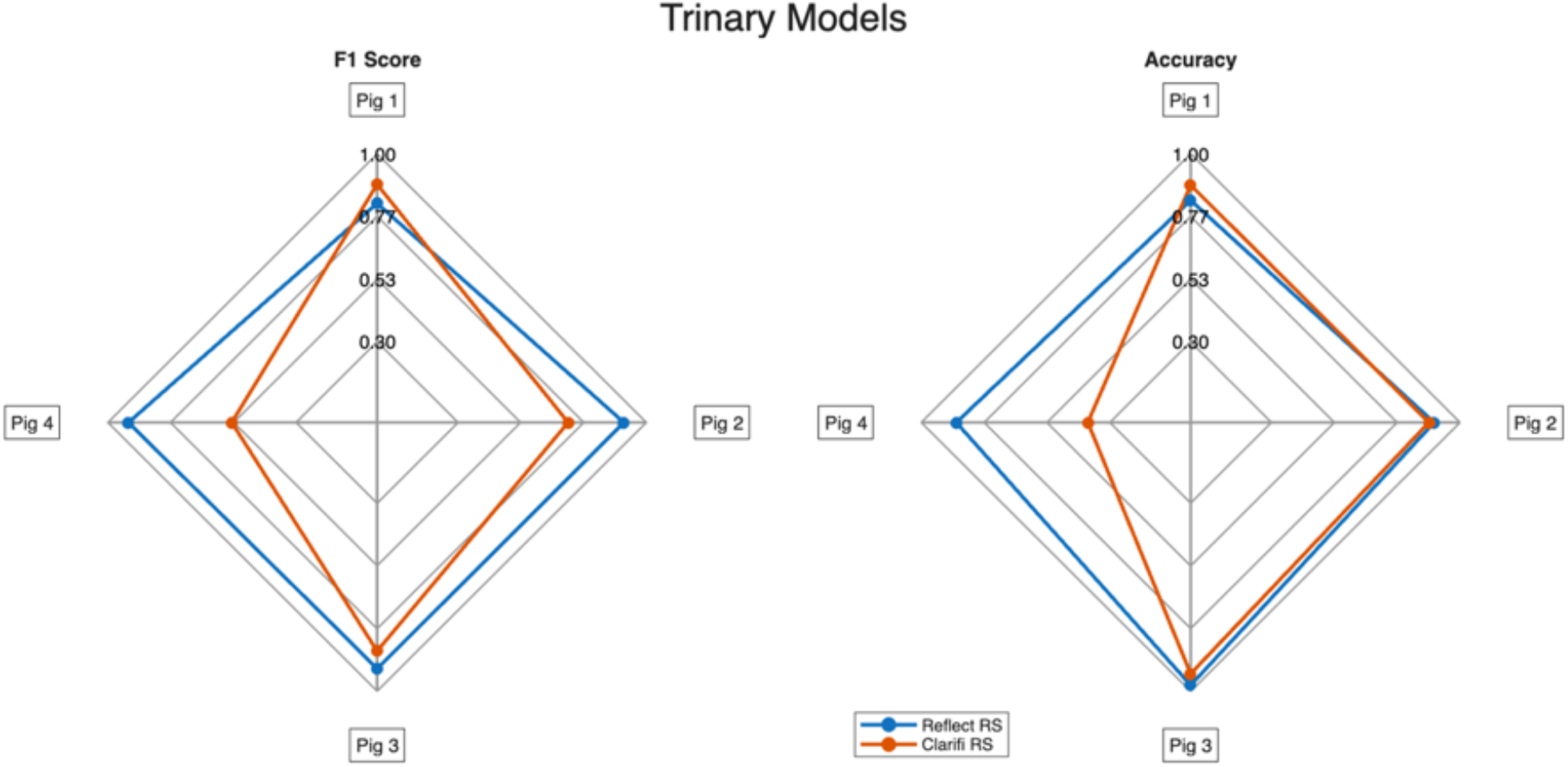
Performance assessment of trinary classifiers. Left: LOSO F1 scores for each pig. Right: LOSO Accuracy in detecting nonviable pixels for each pig. All spider plots were generated using spider_plot() (GitHub: https://github.com/NewGuy012/spider_plot/releases/tag/21.0, accessed July 8, 2026).

Figure 8 shows the appearance of a single, heterogeneous burn, as mapped by the various LOSO-trained models. The top row shows maps produced by the trinary models, while the bottom row demonstrates models trained only on viable/nonviable data, with the unburned skin masked out. Labeled regions are overlaid over all maps, with green representing “viable” and black “nonviable.” The three-category Reflect RS map correctly identifies skin pixels and a majority of “viable” pixels. The Clarifi RS model sometimes fails to distinguish between the three categories. When the “skin” category is removed, both systems classify all “nonviable” pixels and most or all of the “viable” pixels.

**Figure 8.**
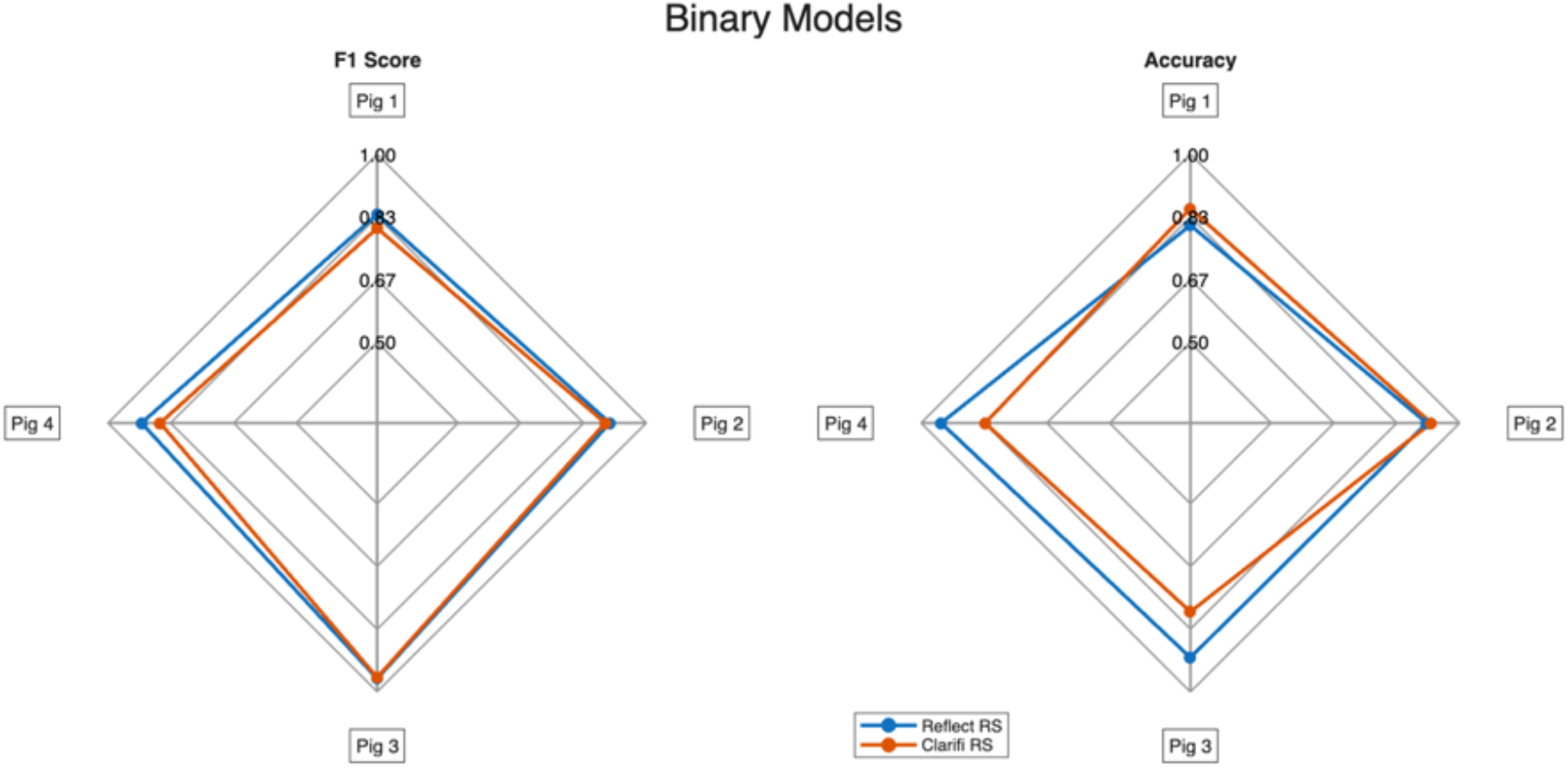
Performance assessment of binary classifiers. Left: LOSO F1 scores for each pig. Right: LOSO Accuracy in detecting nonviable pixels for each pig.

**Figure 8.**
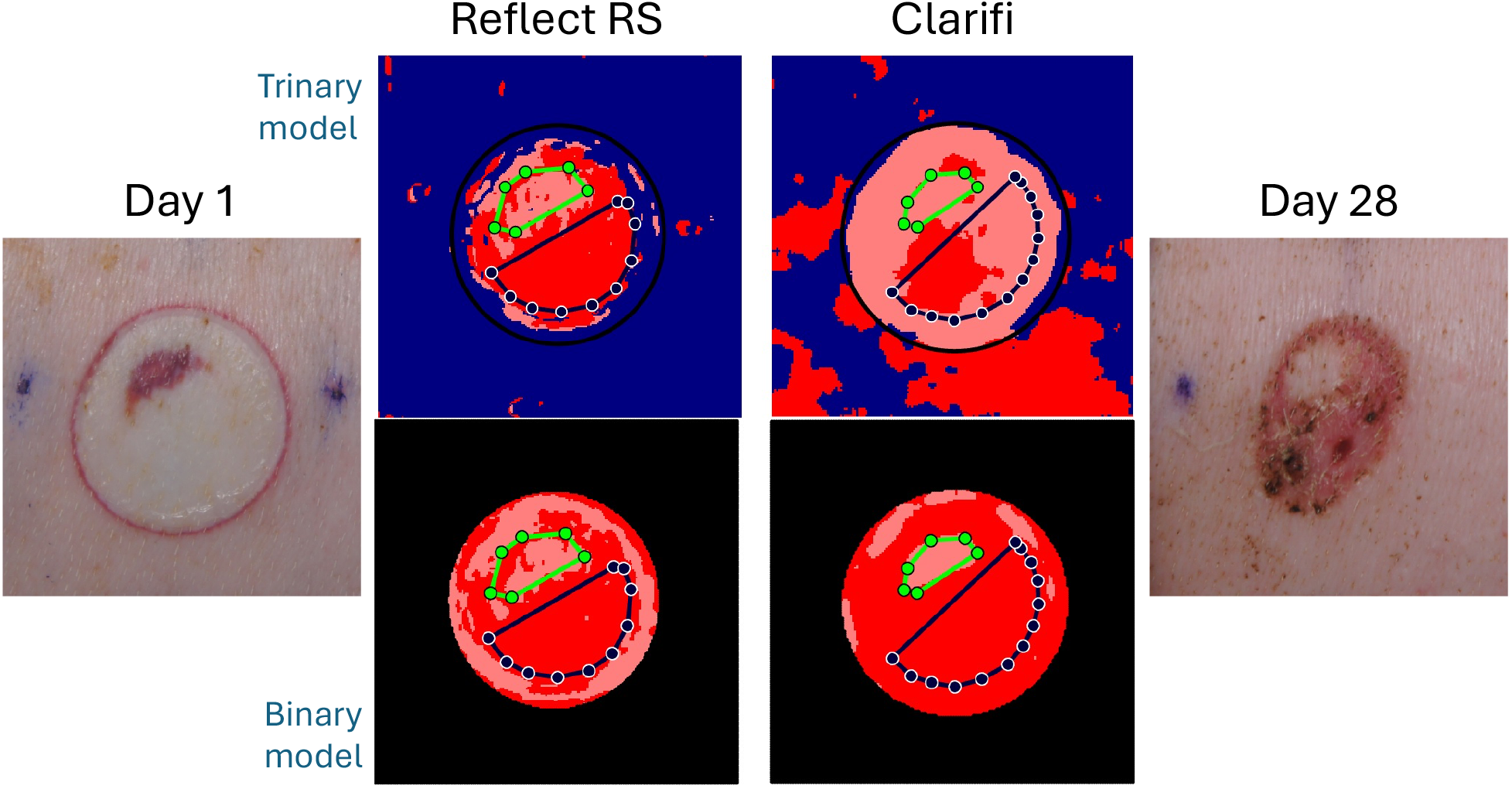
Visual appearance of a classified burn. The top row results from training models on three categories (viable, nonviable, and skin.) The bottom row represents a two-category classification, where the unaffected skin has been masked out by a human, following the outer edge of the hyperperfused burn periphery. To the left and right are color photographs showing the burn’s appearance on Day 1, from which the maps were generated, and Day 28, which was used to create the ROIs used for training and evaluation of the models.

## 4 Discussion

### 4.1 Principal Findings

The primary finding of this study is that a substantially simplified SFDI system retained much of the burn classification capability of a research-grade instrument despite a dramatic reduction in acquired measurement dimensions. Using healing outcome at day 28 as the reference standard, classifiers trained from the full Reflect RS dataset produced the strongest overall performance, achieving mean LOSO F1 scores approaching 0.9. However, classifiers derived from the simplified Clarifi RS dataset maintained strong performance for the clinically relevant task of differentiating viable from nonviable tissue, with mean F1 scores exceeding 0.8. More importantly, the resulting classification maps remained largely consistent with observed healing outcomes and with maps generated using the higher-dimensional Reflect RS dataset (Figures 7 and 8).

These results support the central hypothesis that a clinically practical SFDI system does not necessarily require the extensive spectral and spatial frequency sampling traditionally available in research instruments. While reducing measurement dimensionality predictably decreases performance, the magnitude of that reduction appears manageable within the context of burn severity assessment. This observation has important implications for future translation of SFDI from the research environment into routine clinical and field use.

### 4.2 Impact of Feature Reduction on Burn Classification

One of the motivations for this work was to better understand how much information is required to classify burn severity at early time points. The Reflect RS system acquires measurements across eight wavelengths and multiple spatial frequencies, yielding forty independent reflectance features. In contrast, the Clarifi configuration evaluated here provides only six features that were originally selected for vascular assessment applications rather than burn imaging.

The performance differences observed between the two systems are therefore informative. The Reflect RS consistently outperformed the Clarifi RS in the trinary classification problem (Figure 6), suggesting that additional spectral and depth-sensitive information contributes to improved discrimination among skin, viable burn tissue, and nonviable burn tissue. However, when the task was reduced to the clinically relevant binary distinction between tissue likely to heal and tissue likely to require intervention, the performance gap narrowed considerably (Figure 7). This finding suggests that much of the information contained within the larger Reflect RS dataset may be redundant for the specific purpose of identifying tissue viability.

The qualitative results shown in Figure 8 reinforce this observation. Although the Clarifi RS trinary classifier occasionally struggled to distinguish normal skin from burned tissue, both systems identified the bulk of viable and nonviable regions when evaluated as a binary classification problem. Since the ultimate clinical decision centers on identifying tissue likely to heal versus tissue unlikely to heal by day 28, this distinction is arguably the more important benchmark. The standard of care in burn surgery is that wounds unlikely to heal in this timeframe are autografted to achieve optimal cosmetic and functional outcomes by reducing the likelihood of hypertrophic scar and scar contracture.

### 4.3 Clinical Translation Benefits of Simplified SFDI

An important consideration in any optical imaging system intended for clinical deployment is the tradeoff between information content and operational practicality. Research systems are naturally somewhat optimized for flexibility and discovery, whereas clinical systems must prioritize speed, robustness, ease of use, and workflow integration.

Several characteristics of Clarifi support this translational objective. Compared with Reflect RS, Clarifi acquires fewer images, operates with brighter illumination, and completes image acquisition within only a few seconds. These design choices reduce susceptibility to motion artifacts and improve performance under less controlled imaging conditions. This is particularly important for burn patients, who often experience pain, discomfort, respiratory motion, and other factors that complicate image acquisition.

The value of rapid acquisition becomes especially apparent when considering potential applications beyond specialized burn centers. A portable system capable of providing objective burn severity assessment could be useful in emergency departments, military environments, prolonged field care settings, mass casualty incidents, and resource-limited healthcare systems. Additionally, the handheld portability of such a system would facilitate imaging over larger wound surfaces and anatomic locations, thus improving clinical applicability.

### 4.4 Lessons for Future Burn-Specific Device Design

While the Clarifi RS performed well, the results should not be interpreted as evidence that the current commercial configuration is optimized for burn assessment. The device evaluated here was originally designed for tissue oximetry rather than burn characterization, and its wavelength and spatial frequency selections were presumably chosen accordingly.

The results instead suggest that there is considerable opportunity to develop a burn-specific instrument that occupies a middle ground between Reflect RS and Clarifi. The superior performance of Reflect RS indicates that additional spectral and spatial information contains clinically relevant physiological information. At the same time, the strong performance achieved by Clarifi demonstrates that all forty Reflect RS features are not required.

An important consideration is that SFDI derives information not only from spectral contrast but also from structured illumination. Different spatial frequencies interrogate tissue volumes with different depth sensitivities, allowing the measurement to capture physiological changes occurring beneath the tissue surface. Previous work from our group demonstrated that inclusion of spatial frequency information improves burn classification performance relative to approaches relying solely on planar reflectance measurements. Although Clarifi retained useful predictive capability using a highly reduced feature set, the improved performance of Reflect RS suggests that at least some of the removed spatial-frequency-dependent information contributes meaningful burn severity contrast.

From a translational perspective, the goal should not be to replicate a research-grade instrument in a smaller package. Rather, the goal is to identify the minimum set of wavelengths and spatial frequencies required to make robust clinical decisions. Ultimately, these findings suggest a path toward a lightweight, battery-powered, or handheld SFDI platform capable of providing objective burn severity assessment in environments where conventional imaging systems would be impractical, including emergency response, prolonged field care, and military triage settings.

### 4.5 Importance of Validation Strategy

Previous studies from our group demonstrated that SFDI combined with machine learning can predict burn healing outcomes using calibrated reflectance features acquired from research-grade instrumentation. More recent work using alternative machine learning approaches reached similar conclusions, suggesting that burn classification performance is generally robust across different classifier architectures^15,18^.

An important aspect of the present study is the use of leave-one-subject-out (LOSO) validation. When large numbers of pixels are available from relatively few subjects, validation approaches that mix data from the same subject into both training and testing sets can overestimate real-world performance. LOSO provides a more stringent assessment by training on three animals and evaluating performance on a completely independent fourth animal.

Importantly, the principal findings of this study remained evident under LOSO evaluation. Although performance decreased as measurement dimensionality was reduced, Clarifi-based models continued to provide useful discrimination between tissue that ultimately healed and tissue that progressed to scarring. This suggests that the observed differences between Reflect RS and Clarifi RS primarily reflect differences in available measurement information rather than overfitting or model selection.

### 4.6 Limitations

Several limitations should be acknowledged. First, the study was performed in a controlled porcine model. Although porcine skin remains one of the most widely accepted preclinical models for human burn injury, differences between animal and human wound healing may influence classifier performance^19–21^. Second, ground truth was derived from healing outcomes observed at day 28 rather than histology. This choice of timing for assessing healing outcome may not be optimal; some will argue that non-healing at 14 or 21 days, respectively, are indications for operative intervention and/or autografting.

### 4.7 Summary

These findings demonstrate that substantial reduction in SFDI measurement dimensionality results in only a moderate reduction in predictive performance for early burn severity assessment. Although the research-grade Reflect RS system consistently achieved the highest performance, the simplified Clarifi platform retained the ability to distinguish tissue that ultimately healed from tissue that progressed to scarring.

## 5 Conclusion

This study compared burn severity classification performance between a research-grade SFDI platform (Reflect RS) and a simplified commercial SFDI platform (Clarifi RS) using a porcine model of graded burn injury. As expected, the higher-dimensional Reflect RS dataset produced the strongest overall classification performance. However, despite a substantial reduction in measured features, the Clarifi system retained the ability to distinguish tissue that ultimately healed from tissue that progressed to scarring when evaluated approximately 24 hours following injury.

These findings suggest that clinically useful burn severity assessment does not require the full measurement dimensionality typically available in research-grade SFDI instruments. Rather, a carefully selected subset of spectral and spatial-frequency measurements may be sufficient to support the critical clinical task of distinguishing viable from nonviable tissue.

While the current Clarifi configuration was not designed specifically for burn assessment, the performance observed in this study supports the feasibility of developing compact, rapid-acquisition SFDI instruments optimized for burn care. Such systems have the potential to provide objective decision support in environments ranging from specialized burn centers to emergency response, prolonged field care, and military triage settings.

## Data Availability

The archived version of the data and Matlab code used for this manuscript are available through the Dryad data publishing platform at https://doi.org/10.5061/dryad.0rxwdbsc0.

## Acknowledgements

The work reported in this paper is an expansion of work presented at an SPIE conference^22^. However, that work focused solely on Clarifi RS performance. We thankfully recognize support from the NIH, including the National Institute of General Medical Sciences (NIGMS) (R01GM108634) and the National Institute of Arthritis and Musculoskeletal and Skin Diseases (T32AR080622). We also thank the Arnold and Mabel Beckman Foundation. The content is solely the authors’ responsibility. Any opinions, findings, and conclusions or recommendations expressed in this material are the authors’ and do not necessarily reflect or represent official views of the NIGMS/NIH.

## Author Contribution Statement

CAC analyzed the data, wrote the main manuscript text, and prepared all figures. AM, GTK, and AJD made or suggested substantial edits. AJD, GTK, VJ, and RJC contributed to the animal study design, protocol design/adherence, and funding acquisition. CAC, GTK, AM, TLC, and VJ all participated in carrying out the study. All authors reviewed the manuscript.

## Additional Information

Dr. Durkin is a co-founder of Modulim but does not participate in operation or management of Modulim. He is compliant with UCI and NIH conflict of interest management policy (revisited annually). The other authors have no financial interests or commercial associations representing conflict of interest with the information presented here.

