## Supplemental Figures 1-4 for "Evaluating research-grade and commercial SFDI platforms for burn severity assessment and feature reduction"

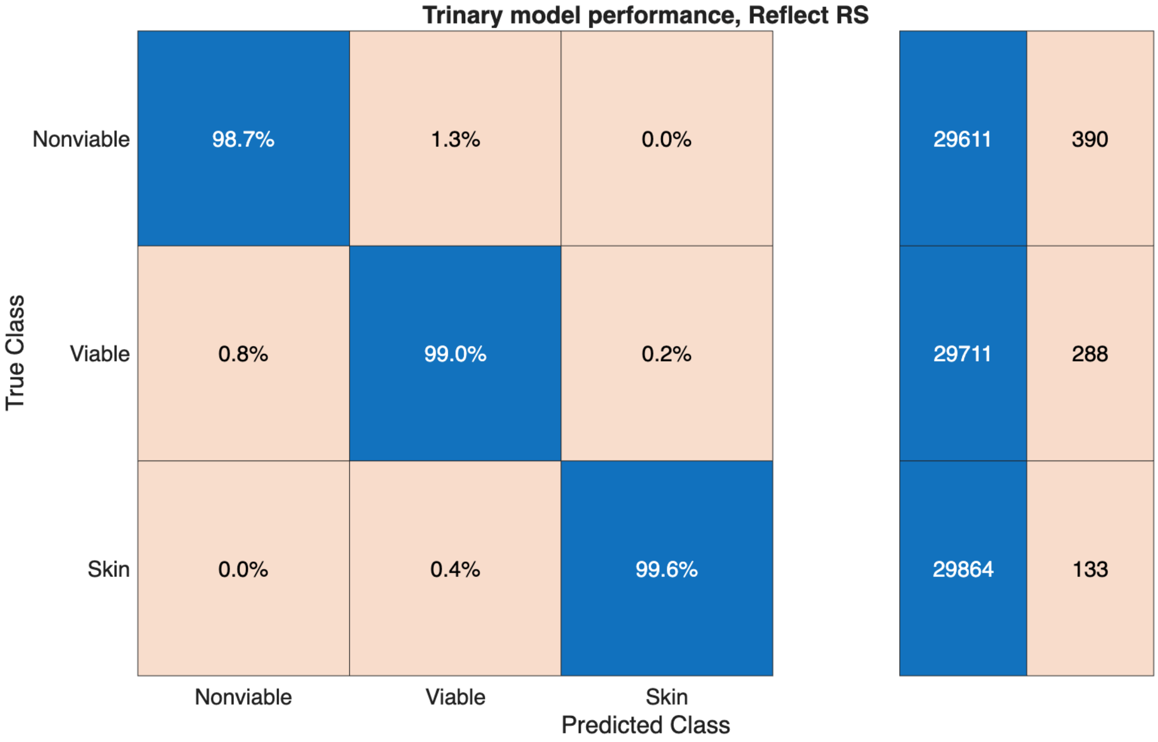


**Supplementary Figure 1:** 10-fold cross-validation matrix for the trinary model trained on Clarifi RS data.

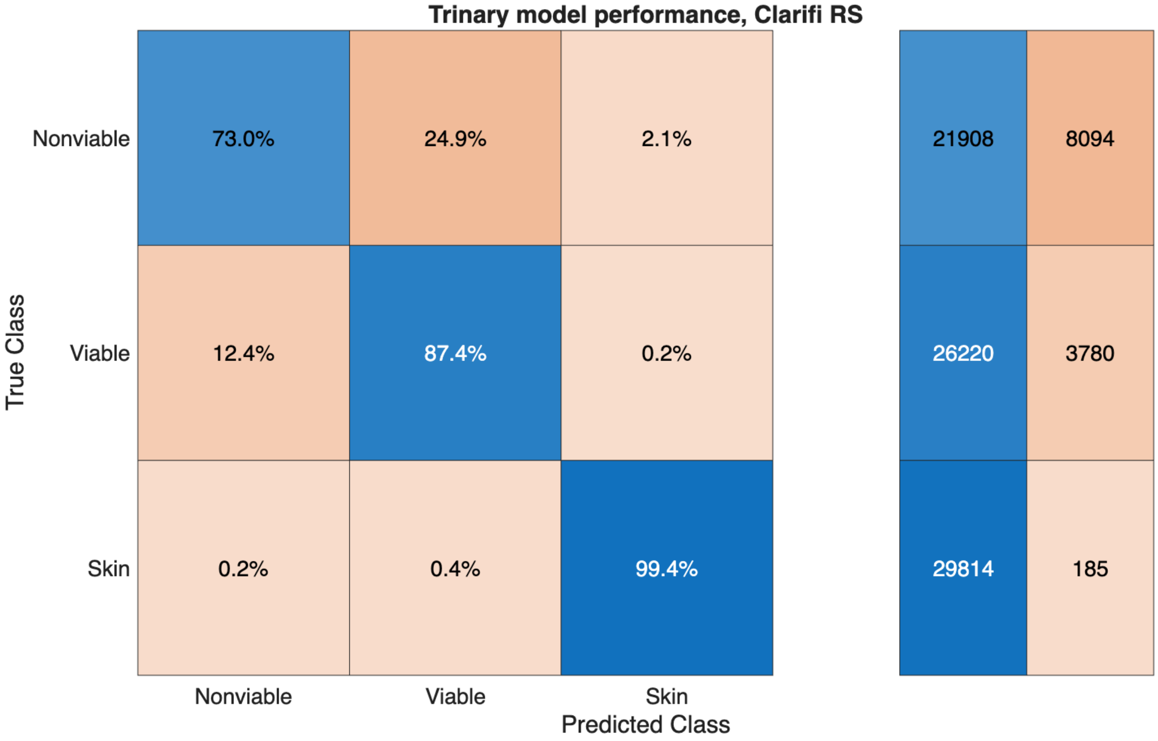


**Supplementary Figure 2:** 10-fold cross-validation matrix for the trinary LOSO model trained on Reflect RS data.

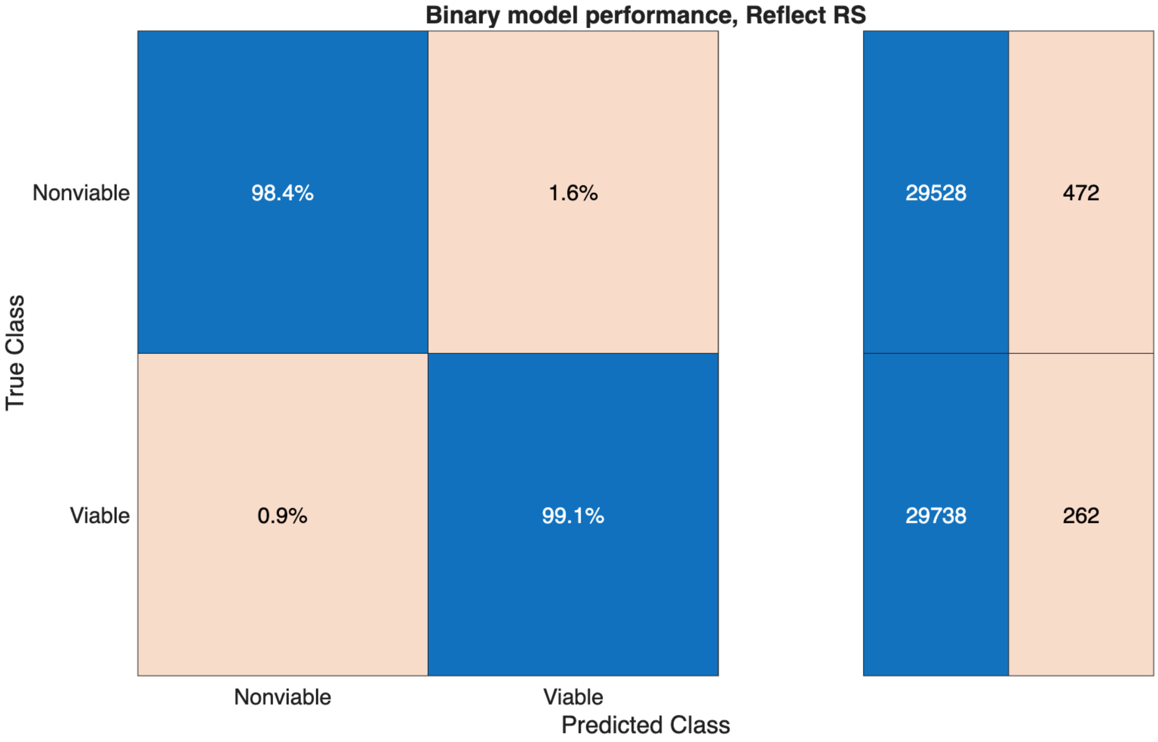


**Supplementary Figure 3:** 10-fold cross-validation matrices for the binary LOSO model trained on Clarifi RS data.

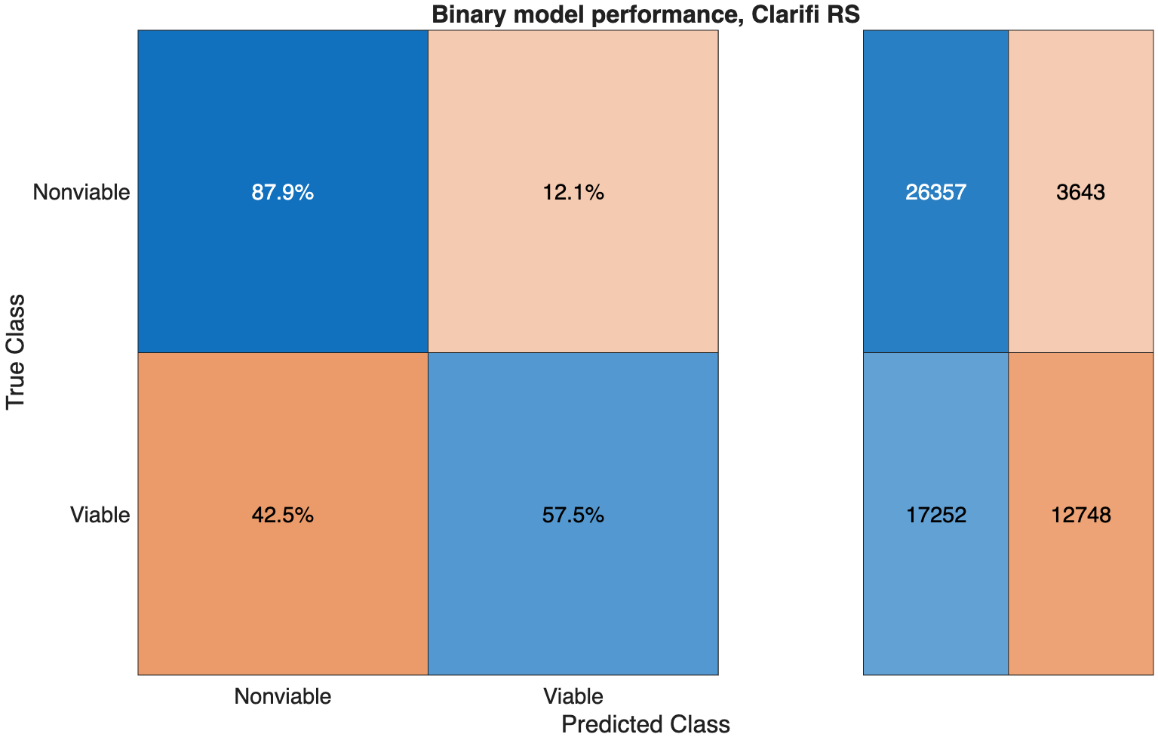


**Supplementary Figure 4:** 10-fold cross-validation matrix for the binary LOSO model trained on Reflect RS data.
